# MACPET: Model-based Analysis for ChIA-PET

**DOI:** 10.1101/272559

**Authors:** Ioannis Vardaxis, Finn Drabløs, Morten B. Rye, Bo Henry Lindqvist

## Abstract

We present Model-based Analysis for ChIA-PET (MACPET) which analyzes paired-end read sequences provided by ChIA-PET for finding binding sites of a protein of interest. MACPET uses information from both tags of each PET and searches for binding sites in a two-dimensional space, while taking into account different noise levels in different genomic regions. MACPET shows favorable results compared to MACS in terms of motif occurrence, spatial resolution and false discovery rate. Significant binding sites discovered by MACPET are involved in a higher number of significant 3D interactions than those discovered by MACS. MACPET is freely available on Bioconductor.

## Background

In the recent years a lot of interest has been placed on understanding the three-dimensional structure of chromosomes inside the cell nucleus [1, 2, 3, 4, 5, 6], as genomes are organized as three-dimensional rather than linear structures in the nucleus of the cell [1]. Those structures play an important role in chromosomal activities such as transcription and regulation of gene expression [1, 2].

The ChIA-PET method allows for analysis of the three-dimensional structure of DNA associated with a protein of interest. It can be used for finding protein binding sites (PBSs) on the genome as well as potential chromatin interactions associated with those proteins. Those interactions provide information on the threedimensional genome structure [6].

ChIA-PET data contain short DNA sequences ~ 20 base pairs (bp), which are called tags. Each tag is ligated to a half-linker sequence, either A or B (often TAAG for linker A and ATGT for linker B). Each of those half-linkers also contain the site for the restriction enzyme used to cut the sequences for releasing the tags. For example GTTGGA for the Mmel restriction enzyme which cuts 20 bp from its restriction site to reveal the 20 bp long tag sequence. Pairs of tag-half-linker products are connected to each other by proximity ligation to form tag-linker-tag products named paired-end-tags (PETs). The final linker sequence re-veals different combinations of the two half-linkers A and B [7].

The combination of the half-linkers classifies the PETs into three categories. *Ambiguous* PETs are those for which any of their half-linkers is missing. *Chimeric* PETs are those with half-linkers A/B or B/A and are derived from random ligations between different ChIP complexes. Finally, *non-chimeric* PETs are those with half-linkers A/A or B/B. Only the non-chimeric PETs are considered for the ChIA-PET analysis [7].

After classifying the PETs based on their half-linkers, the linker sequences are removed from the non-chimeric PETs [7], and the tags of each PET are separately mapped on the genome [8, 9]. The location to which the tags are mapped classifies the PETs into three categories (see figure 1) [7, 10]. *Self-ligated* PETs are products of self-circularization ligation of a single DNA fragment. They consist of tags which belong to the same chromosome and strand, and which have same orientation and short genomic span between them. Furthermore, each PET has the same chance of being sequenced on both strands which will result in both tags being mapped either on Watson or on Crick strand. *Intra-chromosomal* PETs consist of tags which belong to the same chromosome, have long genomic distance between them, and may have been mapped on different strands or with different orientation. *Inter-chromosomal* are PETs with the same characteristics as the intra-chromosomal ones, but their tags are mapped on different chromosomes. Intra- and inter-chromosomal PETs correspond to two DNA fragments from different genomic regions bound to the same protein of interest and ligated to each other during the ligation process [7, 10].

**Figure 1.**
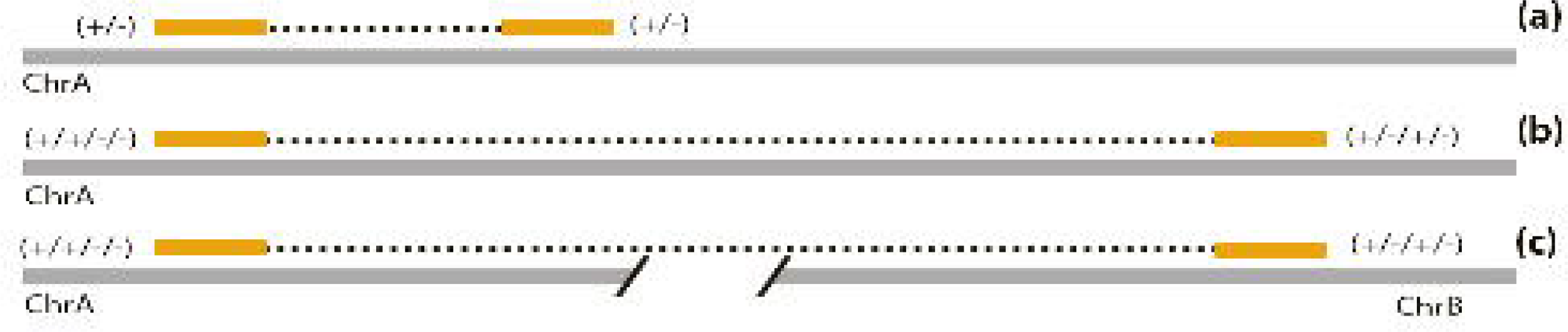
Illustration of PET types. (a) Self-ligated PETs with both tags on same chromosome and strand, and short genomic distance between them. (b) Intra-chromosomal PETs with tags on same chromosome, with any strand combination and long genomic distance between them. (c) Inter-chromosomal PETs with tags on different chromosomes and with any strand combination.

The process of creating ChIA-PET data is composed of many experimental steps, each of which may introduce some noise to the data [8, 11, 7]. Noise PETs are usually defined as those that show a random positioning on the genome without creating significant peaks [7]. However, noise might not be evenly distributed across the genome, but might gather around certain regions [11]. This suggests different amounts of noise in different regions. Furthermore, noise can also be generated by PCR clonal amplification. These PETs might show significant overlap and be misclassified as PBSs by peak-calling algorithms. For reducing that kind of noise, Li et al. [7] proposes merging all PETs for which both of their tags overlap ±1 bp with another PET’s tags.

Self-ligated PETs are used for identifying PBSs by finding significant peaks of overlapping PETs on the genome. The tags of those PETs would pile up creating two peaks, one upstream and one downstream from the true PBS location, where the exact binding position exists somewhere between the two peaks [7, 10].

MACS is a widely used algorithm for finding PBSs on DNA for ChIP-Seq data using a non-parametric model [12]. However MACS is also used for ChIA-PET data by using only the 5-end tag of each PET. Because PETs can be sequenced from either strand with the same probability, using only the 5-end tag of each PET would reveal an upstream and a downstream density of the Watson and Crick tags, respectively, with the PBS location positioned in the middle. MACS identifies and separates the two densities around each PBS by scanning the genome with a user-specified window. It then shifts the two densities towards each other to find the precise binding location. Finally, it merges candidate PBSs which overlap for creating a single one [12]. However, if the window is too large the PBSs might be overestimated, and if it is too small the probability of false positives increases [11]. The choice of the window size might be a challenge for the user.

It would be reasonable to assume that using both tags of the self-ligated PETs provided by ChIA-PET would result in better identification of the PBSs. This is because if a PET belongs to a PBS, then both of its tags should be mapped around that PBS, irrespective of strand. Additionally, using a parametric model which takes into account specific characteristics of the PBSs for identifying their location, might be more efficient than a non-parametric. To be more specific, it might be reasonable to expect that the distribution of the upstream peak of a PBS would be negative-skewed towards the PBS and with longer left tail, since more upstream tags will be mapped on the left side of the PBS. Accordingly, the distribution of the downstream peak would be positive-skewed towards the PBS and with longer right tail, since more of the downstream tags will be mapped on the right side of the PBS. Predicting more accurate PBS locations should result in better discovery of significant interactions between those PBSs.

Intra- and inter-chromosomal PETs are used for finding interactions between PBSs which are previously identified by the self-ligated PETs. Those interactions provide information about the three-dimensional structure of the genome and how it is folded in the nucleus of the cell [7, 10].

MANGO is a complete ChIA-PET pipeline which uses MACS for finding significant PBSs and searches for significant interactions between those PBSs by taking the distance between them into account. Moreover, the user can choose which stage of the MANGO analysis to run, as well as provide PBSs found by algorithms other than MACS. MANGO has been proven to give more accurate results than other algorithms of the same kind [13].

In the current paper we present Model-based Analysis for ChIA-PET (MACPET), an efficient method for discovering PBSs using ChIA-PET data. MACPET uses both tags of each self-ligated PET and estimates the PBSs using two-dimensional parametric mixture models. Modeling the self-ligated PETs in two dimensions, one dimension for each tag, and representing them as dots in a two-dimensional space, ensures that in order for a self-ligated PET to belong to a PBS, both of its tags need to belong to it. MACPET identifies the upstream and downstream peaks of each PBS by taking into account potential skewness of the peaks. Since both tags of each self-ligated PET are used, MACPET does not use strand information of the tags. Furthermore, MACPET models non-overlapping genomic regions separately and evaluates noise locally, which results in better identification of noise PETs and excludes the need for user-specified values. Finally, MACPET also implements the preliminary stages of ChIA-PET analysis like linker identification, linker trimming, mapping to the reference genome and PET classification. The output of MACPET can be directly used in the MANGO algorithm. MACPET is publicly available at Bioconductor. It is mainly implemented in C++ and is thus fast and supports all relevant platforms.

## Results

We compare MACPET with MACS on six ChIA-PET datasets publicly available at NCBI [14]. Table 1 presents the datasets used and shows the results of the first three stages of MACPET analysis. Because MACS cannot filter, trim, map or classify the PETs we used MACPET for those stages. The self-ligated PETs found by MACPET are then used in MACS for binding site analysis. In the main text we only present results from the ESR1 (MCF-7), CTCF (MCF-7) and CTCF (K562) datasets. The results of the remaining datasets can be found in the supplementary material available online. However, we will refer to them in the main text.

**Table 1.** Description of the datasets.

| Name | GEO | PETs | Ambiguous | Chimeric | NN | Usable | Final PETs | Inter-chrom. | Intra-chrom. | Self-ligated |
| --- | --- | --- | --- | --- | --- | --- | --- | --- | --- | --- |
| ESR1 (MCF-7) | GSM970212 | 26170257 | 165093 | 2492122 | 620598 | 22892444 | 6575793 | 1765891 | 164974 | 534284 |
| CTCF (MCF-7) | GSM970215 | 119959634 | 7105105 | 11018238 | 317231 | 101519060 | 53526399 | 20630265 | 2729210 | 5872607 |
| CTCF (K562) | GSM970216 | 195436387 | 42363105 | 9477776 | 1893878 | 141701628 | 80927884 | 2824894 | 1531945 | 2337518 |
| H3K4me1 (K562) | GSM1436263 | 162190720 | 23144511 | 30577815 | 465092 | 108003302 | 38161523 | 31963485 | 2483809 | 2867067 |
| H3K27ac (K562) | GSM1436262 | 165109173 | 32418202 | 53756946 | 683759 | 78250266 | 25168926 | 19265500 | 1770107 | 3167743 |
| POL2 (K562) | GSM970213 | 37889691 | 3920503 | 409894 | 453521 | 33105773 | 17473165 | 2919406 | 4068174 | 7387093 |

Name of the datasets (Name), GEO number of the datasets (GEO) and total number of initial PETs of the datasets (PETs). Total PETs classified as Ambiguous, Chimeric, Usable (non-chimeric), as well as the total PETs with non-standard residues (NN). Total valid paired and mapped PETs used (Final PETs), as well as their classification into inter-chromosomal, intra-chromosomal and self-ligated.

Figure 2 shows the self-ligated and intra-chromosomal separation cut-off of the three datasets (for the other three datasets see figure S2 in supplementary material available online). The self-ligated data was then used in both MACPET and MACS for finding significant PBSs on each dataset. For both MACPET and MACS we declared significant PBSs with false discovery rate (FDR) cut-off at 0.05, mainly because this is the default cut-off for PBSs which are used in the interaction analysis algorithm as we discuss later.

**Figure 2.**
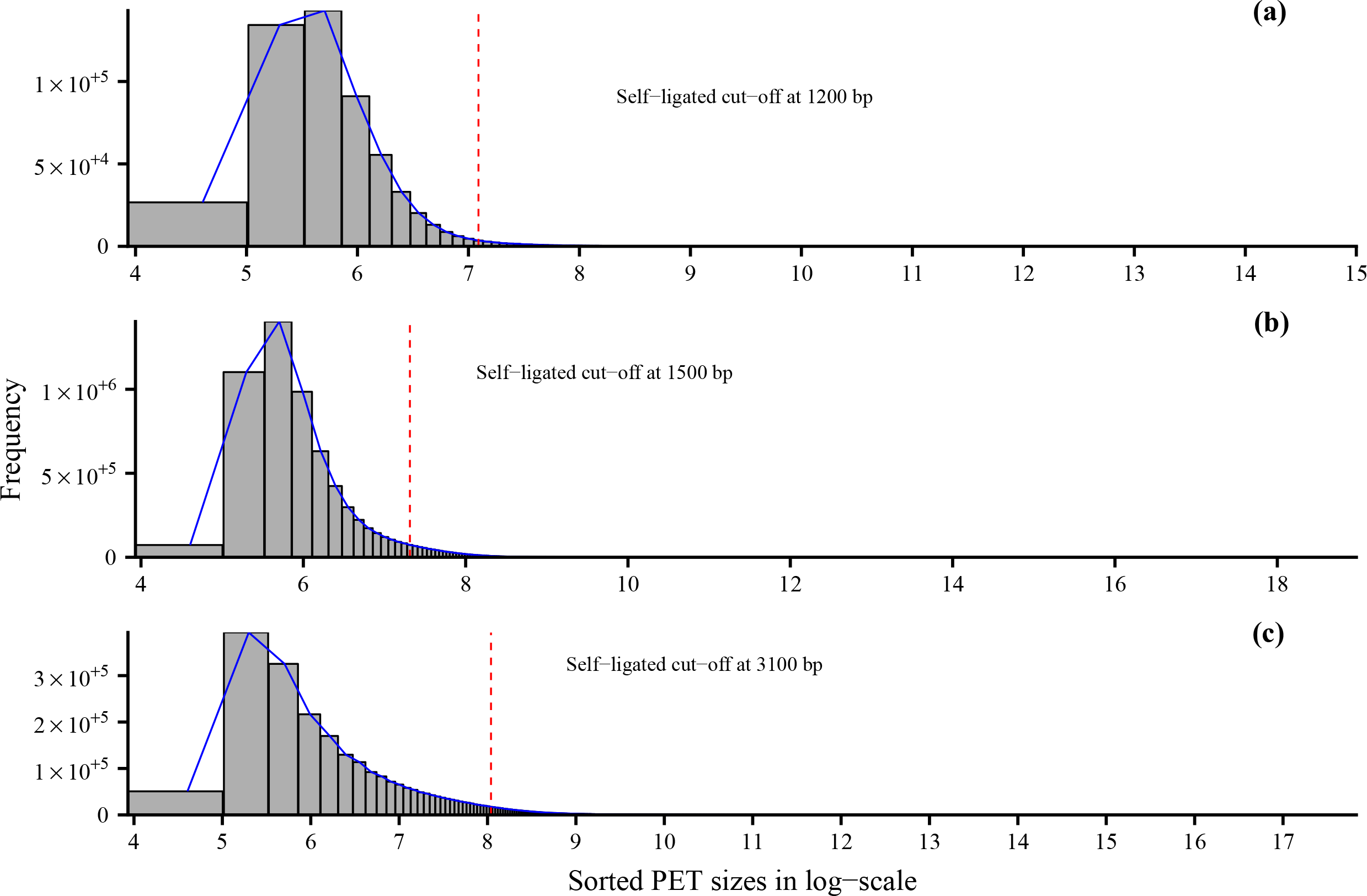
Self-Intra cut-off. Self-ligated and Intra-chromosomal cut-off for the three datasets. (a) ESR1 (MCF-7), (b) CTCF (MCF-7), (c) CTCF (K562). The x-axis are the lengths of the PETs in log scale, while the y-axis is the frequency. The dashed line represents the cut-off point, where the self-ligated PETs are on its left side and the intra-chromosomal on its right.

For the ESR1 (MCF-7), CTCF (MCF-7) and CTCF (K562) datasets we can investigate the association of the significant PBSs with the expected motifs (ESR1 and CTCT accordingly). Using 200 bp windows centered at the precise PBS locations for the top 5000 most significant PBSs from MACPET and MACS, we compare the quality and precision of those bindings in terms of motif occurrence (the percentage of PBSs associated with the expected motif) and spatial resolution (the distance of the PBS location to the expected motif). For doing so, we used the rGADEM algorithm [15] for de novo motif analysis, and then the MotIV algorithm [16], for keeping only the most common motif on each dataset. rGADEM applies a stochastic algorithm for de novo motif discovery and it is therefore not guaranteed to give identical results on each run [17]. Therefore, we run rGADEM five times for both MACPET and MACS (using MotIV after each run) and we take the mean among the runs as the final result. For both MACPET and MACS the most common motifs for each run were the expected motif for each dataset.

Figures 3 (a-c) show the motif occurrence for each dataset. MACPET results in a higher number of PBSs associated with the expected motif than those from MACS, for each dataset. Figures 3 (d-f) show the spatial resolution of the PBSs, where only PBSs with distance less than 50 bp from the expected motif are taken into account. The locations of the MACPET PBSs are more precise as they are closer to the expected motif location than those of MACS.

**Figure 3.**
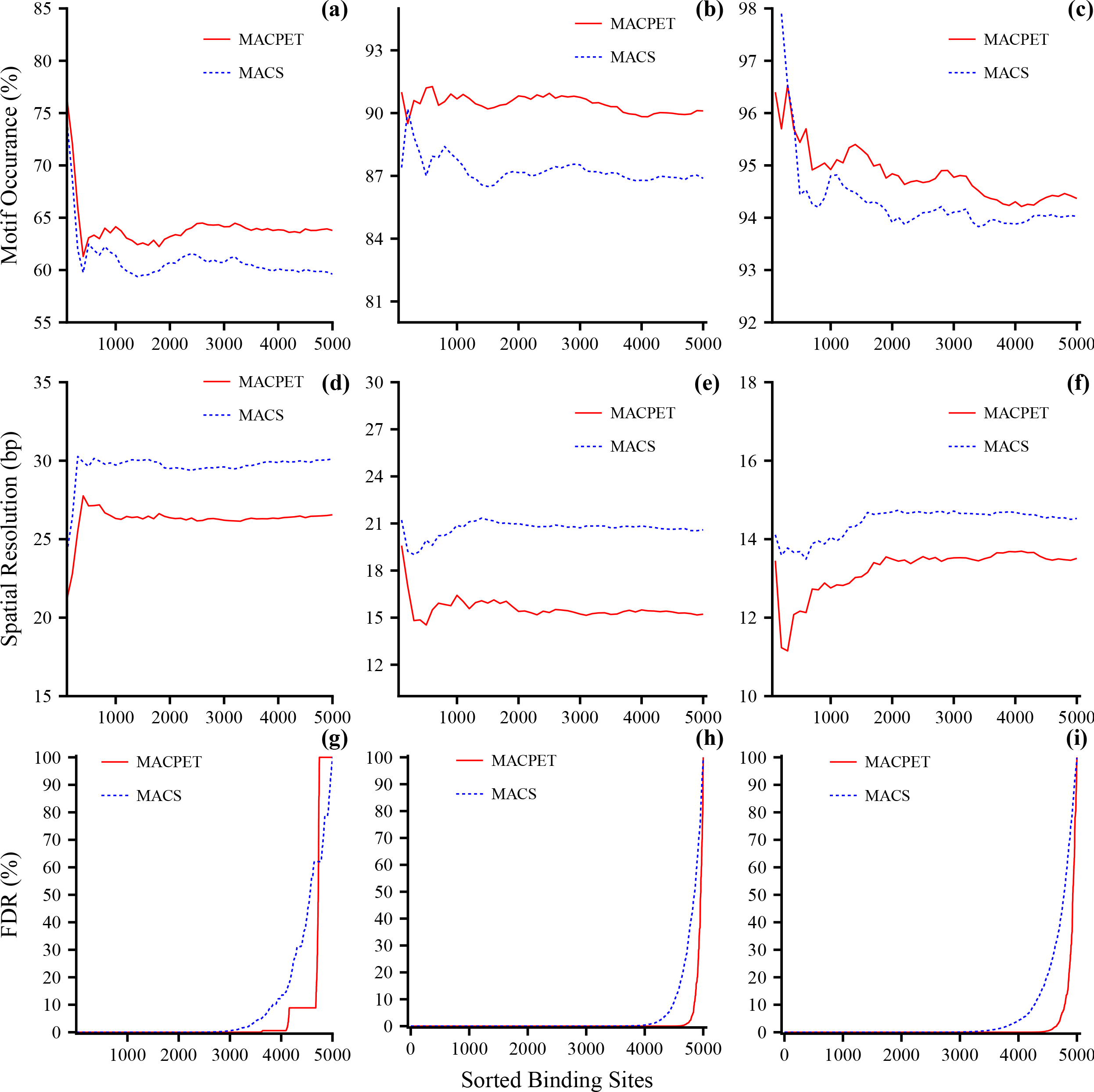
De novo motif discovery and FDR. Comparison of motif discovery, spatial resolution and FDR between MACPET and MACS. The x-axis for all plots is the top 5000 PBSs, sorted by significance in descending order. (a-c) Motif occurrence (y-axis) for (a) ESR1 (MCF-7), (b) CTCF (MCF-7), (c) CTCF (K562). (d-f) Spatial resolution (y-axis) for (d) ESR1 (MCF-7), (e) CTCF (MCF-7), (f) CTCF (K562). (g-i) FDR (y-axis) for (g) ESR1 (MCF-7), (h) CTCF (MCF-7), (i) CTCF (K562).

For evaluating the inference methods of MACPET and MACS, we compare the FDR of the first 5000 most significant PBSs. Figures 3 (g-i) show the FDR for the three datasets (for the other three datasets see figures S3 (a-c) in supplementary material available online). MACPET results in lower FDR than MACS in general for all six datasets, which is an indication that fewer false PBSs will be found by MACPET.

Figures 4 (a-c) show Venn diagrams for the significant PBSs which are in common for MACPET and MACS, for the three datasets (for the rest of the datasets see figures S4 (a-c) in supplementary material available online). There are in general many PBSs which are common for the two algorithms. However, it is noticeable that MACS finds far more significant PBSs than MACPET. In figures 4 (d-f), on the other hand, one can see that MACPET finds stronger PBSs in terms of total tags than MACS for all six datasets (for the rest of the datasets see figures S4 (d-f) in supplementary material available online).

**Figure 4.**
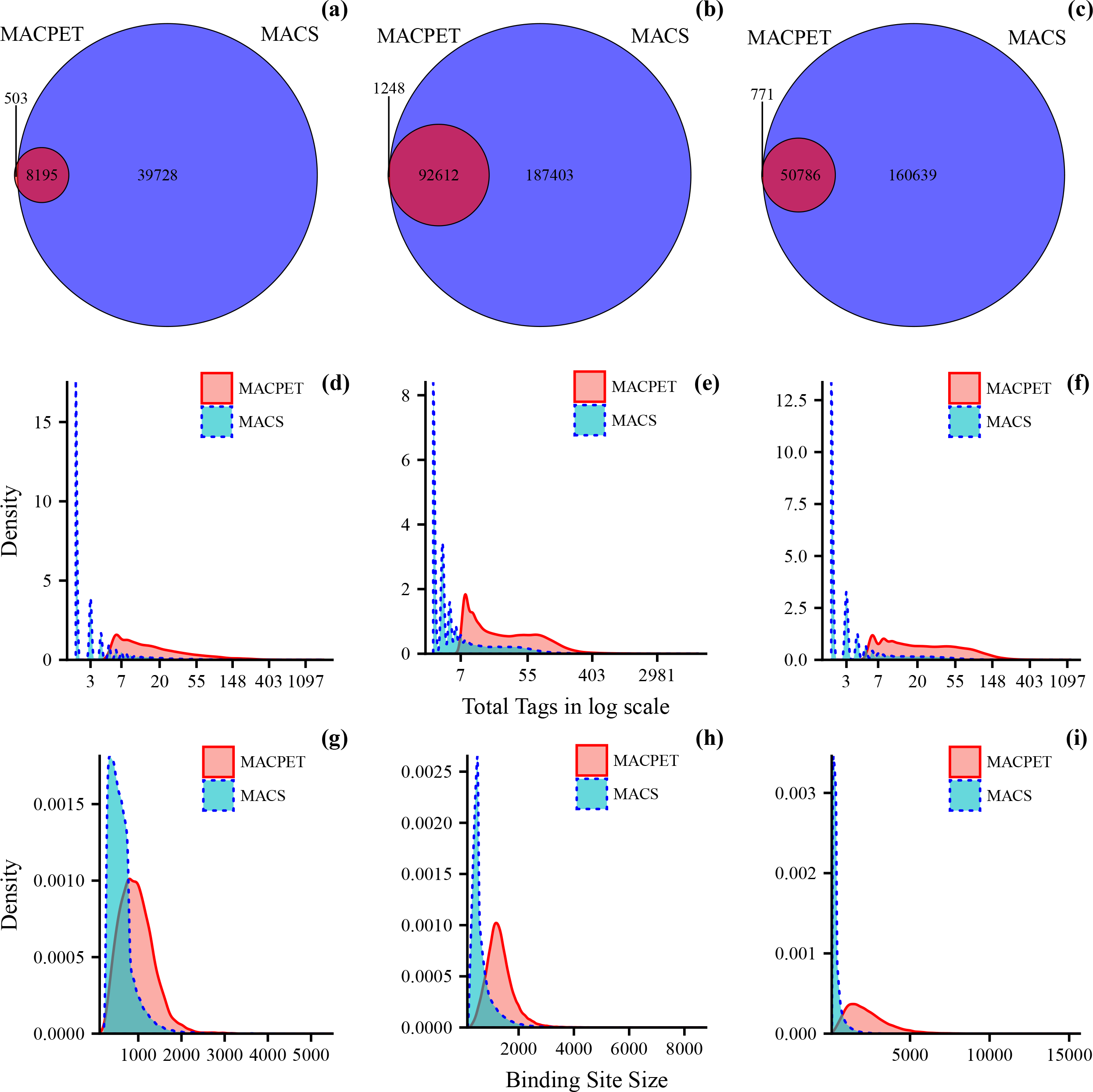
Comparison of significant binding sites. (a-c) Venn diagrams of the significant PBSs for MACPET and MACS for ESR1 (MCF-7)(a), CTCF (MCF-7)(b), CTCF (K562)(c). (d-f) densities for the total number of tags in each significant PBSs from MACPET and MACS for (d) ESR1 (MCF-7), (e) CTCF (MCF-7), (f) CTCF (K562). The x-axis is the total tags in log scale and the y-axis is the density of the total tags. (g-i) densities for the sizes of the significant PBSs from MACPET and MACS for (g) ESR1 (MCF-7), (h) CTCF (MCF-7), (i) CTCF (K562). The x-axis is the sizes of the significant PBSs and the y-axis is their density.

Additionally, comparing the interval sizes of the PBSs in figures 4 (g-i), we can see that MACPET seems to result in larger, but probably more realistic PBS intervals than MACS (for the rest of the datasets see figures S4 (g-i) in supplementary material available online).

Moreover, we used the fifth stage in MANGO algorithm for investigating the potential benefits of the PBSs from MACPET over those from MACS in terms of interaction analysis and 3D DNA structure. We used the significant PBSs found by MACPET and MACS as inputs in MANGO (FDR< 0.05 which is the default for peak-calling in MANGO), as well as the intra-and inter-chromosomal PETs classified by MACPET.

MANGO gives the option to extend the PBS intervals in both sides with a user specified window (500 bp being the default) [13]. Because MANGO merges the extended PBSs before running interaction analysis [13], we run MANGO on a sequence of extending windows 0,100,…, 900,1000. This allows us to investigate how different extending windows affect the merging of the PBSs and thus the interactions. The rest of MANGO parameters are kept at default for both MACPET and MACS.

Figures 5 (a-c) show the total number of significant interactions (FDR< 0.05) for each extension window for MACPET and MACS (for the rest of the datasets see figures S5 (a-c) in supplementary material available online). MACPET gives higher total number of significant interactions than MACS for all six datasets and all windows. In figures 5 (d-f) we also compared the total PBSs involved in interactions for MACPET and MACS (for the rest of the datasets see figures S5 (d-f) in supplementary material available online). PBSs found by MACPET are more involved in interactions than those from MACS.

**Figure 5.**
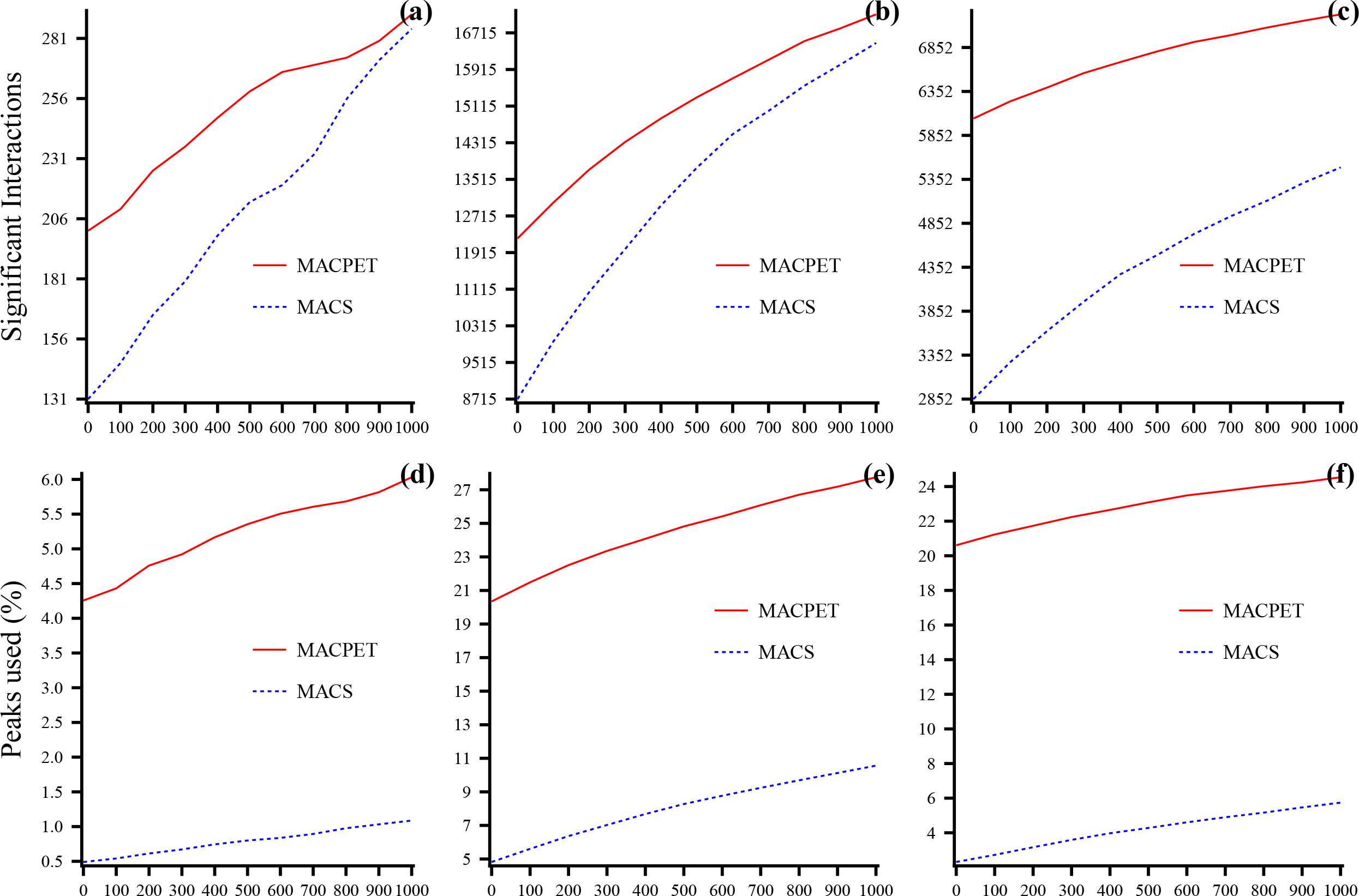
Comparison for MANGO interactions. Comparison of MANGO interaction results between significant PBSs from MACPET and MACS, for different PBSs extension windows. For all the plots, the x-axis is the number of bp. Each PBS interval was extended from either side before running MANGO. (a-c) total significant interactions (y-axis) for (a) ESR1 (MCF-7), (b) CTCF (MCF-7), (c) CTCF (K562). (d-f) proportion of significant PBSs involved in significant interactions (y-axis) for (d) ESR1 (MCF-7), (e) CTCF (MCF-7), (f) CTCF (K562).

Finally, we consider only interactions for the 500 bp extension window. Figures 6 (a-c) show the Venn diagrams for the common interactions between MACPET and MACS (see figures S6 (a-c) for the rest of the data in supplementary material available online). In general there are many common interactions between MACPET and MACS. However MACPET reveals many more interactions than MACS. Figures 6 (d-f) show the distance of the interactions (see figures S6 (d-f) for the rest of the datasets in supplementary material available online), where MACPET seems to result in slightly longer interactions for all the datasets.

**Figure 6.**
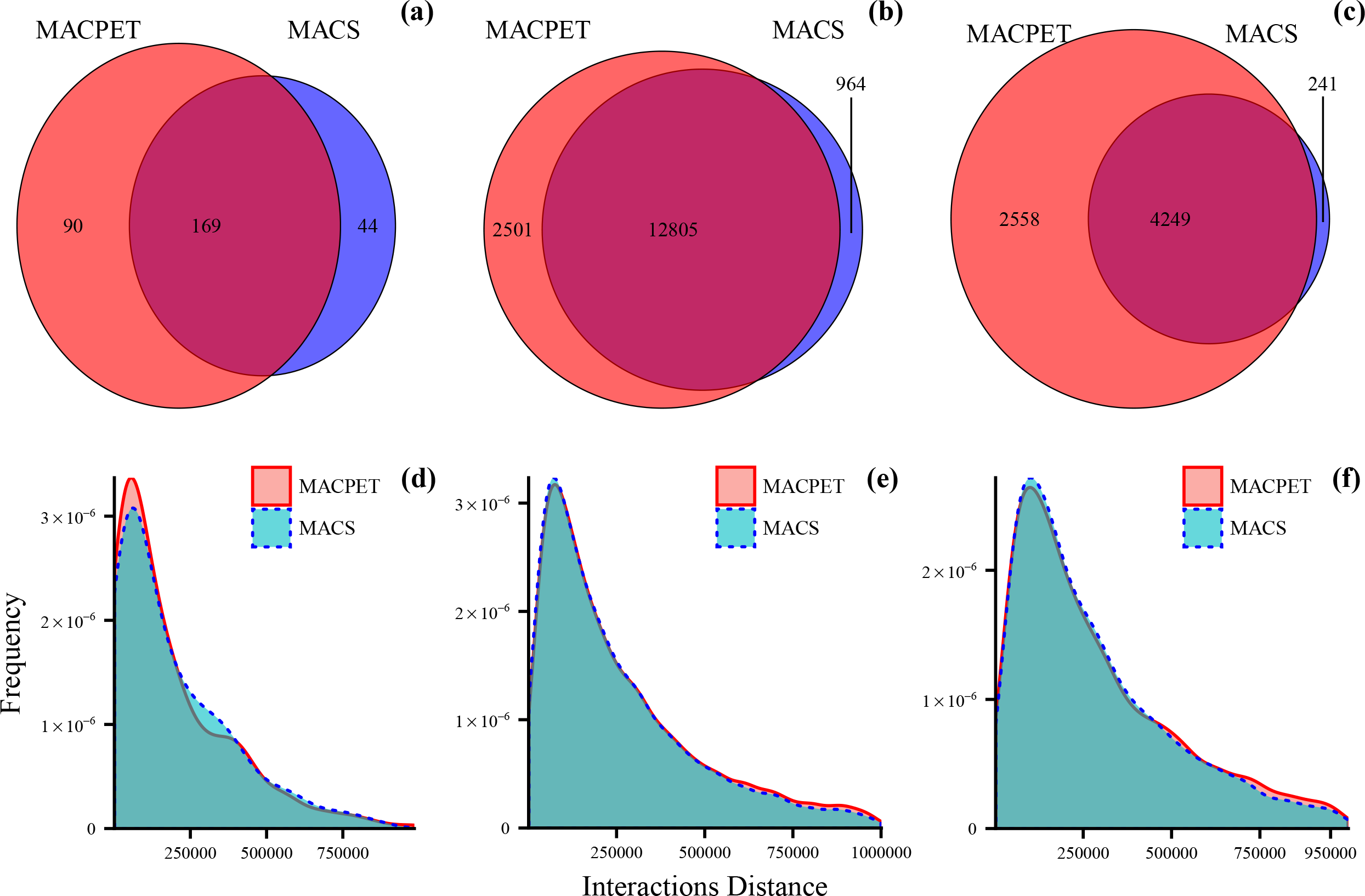
Comparison for MANGO interactions of 500 bp window extension. (a-c) Venn diagrams from significant interactions for a 500 bp extension window for the significant PBSs from MACPET and MACS for (a) ESR1 (MCF-7), (b) CTCF (MCF-7), (c) CTCF (K562). (d-f) density plots forthe distances of the significant intra-chromosomal interactions from MACPET and MACS for (d) ESR1 (MCF-7), (e) CTCF (MCF-7), (f) CTCF (K562). The x-axis is the sizes of the intra-chromosomal interactions and the y-axis is their density.

## Discussion

We compared MACPET with MACS since the latter is one of the most used algorithms for discovering PBS. PICS [18] is another known algorithm for binding site analysis which also uses only the 5-end tag when used for ChIA-PET data. However, PICS needs control data for computing the FDR for the PBSs, and control data were unavailable for the datasets used in the analysis. Without an FDR estimate it was not possible to subset the most significant PBSs from PICS and thus, we could not compare PICS with MACPET.

We showed that MACPET discovers fewer significant PBSs than MACS. This is expected since MACPET models noise locally using mixture models, which results in weaker overlapping PETs being categorized as noise, while MACS categorizes them as PBSs. This is consistent with the PBSs’ FDR for MACPET and MACS. The FDR of MACPET PBSs is much lower than those for MACS, indicating that many of the PBSs found by MACS are actually mistaken as PBSs, but they should rather be classified as noise (i.e., false positives). This result was also consistent with the strength of the PBSs from MACPET and MACS, where we showed that PBSs found by MACPET contain a higher number of tags than PBSs found by MACS. Which is expected since MACPET forces both tags of each self-ligated PET to be part of a PBS.

Moreover, we showed that MACPET results in better identification or PBSs as well as more accurate positioning of PBSs than MACS. Although MACPET finds fewer significant PBSs, those PBSs are associated with the expected motif in higher frequency than those from MACS. This indicates once more that MACS have discovered a higher number of false PBSs with no motif association. Moreover, PBSs found by MACPET were closer to the exact motif position than those from MACS. Consequently, this confirms the assumption that using both tags of each PET in ChIA-PET data results in more accurate PBS locations.

Additionally, we showed that MACPET results in PBSs with longer and, overall, more flexible intervals. The skewness that MACPET implements when modeling PBSs seems to reflect the characteristics of the proteins being modeled. For example, PBSs from the datasets ESR1 (MCF-7), CTCF (MCF-7) and CTCF (K562), which are transcription factor proteins and it is known that they bind at specific location, give smaller intervals than the dataset POL2 (K562), which is a polymerace protein and it is known to bind in wide locations. MACS also captures the characteristics of the proteins, however not as much as MACPET does, because MACS’ model is not as flexible.

We also investigated how PBSs found by MACPET affect the 3D genome interactions, compared to those from MACS. We used the significant PBSs found by MACPET and MACS in the interaction stage of the MANGO algorithm. We showed that, although the total number of PBSs given as input in MANGO was much higher for MACS than for MACPET, MACPET resulted in a higher number of significant interactions between its PBSs irrespective of the extending window. Furthermore, a higher portion of PBSs found by MACPET are involved in interactions than those from MACS. This also indicates that the quality and precision of the PBSs found by MACPET are better than those from MACS.

Finally, we showed that PBS found by MAPCET are involved in slightly longer interactions than PBS found by MACS. It is well known that PBS which are close to each other in genomic distance, tend to randomly interact more often than PBS which are separated by long genomic distance [3]. This also indicates that the PBS found by MACPET are more accurate than those found by MACS.

## Conclusions

The aim of the study was to create an algorithm-pipeline which would take advantage of all the available information provided by paired-end data like ChIA-PET for discovering PBSs. The reason behind this was that identifying more accurate PBS locations should result in more robust identification of interactions. As intra-and inter-chromosomal PETs connect PBSs by being mapped near the PBSs’ binding locations, improperly identified PBSs might result in weak or even inaccurate interactions. We created MACPET, which runs a ChIA-PET data analysis including stages for linker trimming, mapping to the reference genome, PET classification, as well as a new statistical method for discovering PBSs using both tags of each PET. We showed that using all the available information from the paired-end data, combined with a more flexible model when discovering PBSs, is very important and leads to the discovery of a higher number of interactions between those PBSs. Those interactions might reveal new insights of the 3D DNA structure which might not have been found if using only the one tag of the paired-end data for finding PBSs. Finally, although the output from MACPET can be directly usedin MANGO for interaction analysis, we are planning to implement a new interaction model in MACPET in the near future.

## Methods

MACPET currently implements a four-stage analysis of ChIA-PET data. Each of those stages (0-3) is briefly discussed in the following Sections. Figure 7 shows a complete MACPET pipeline.

**Figure 7.**
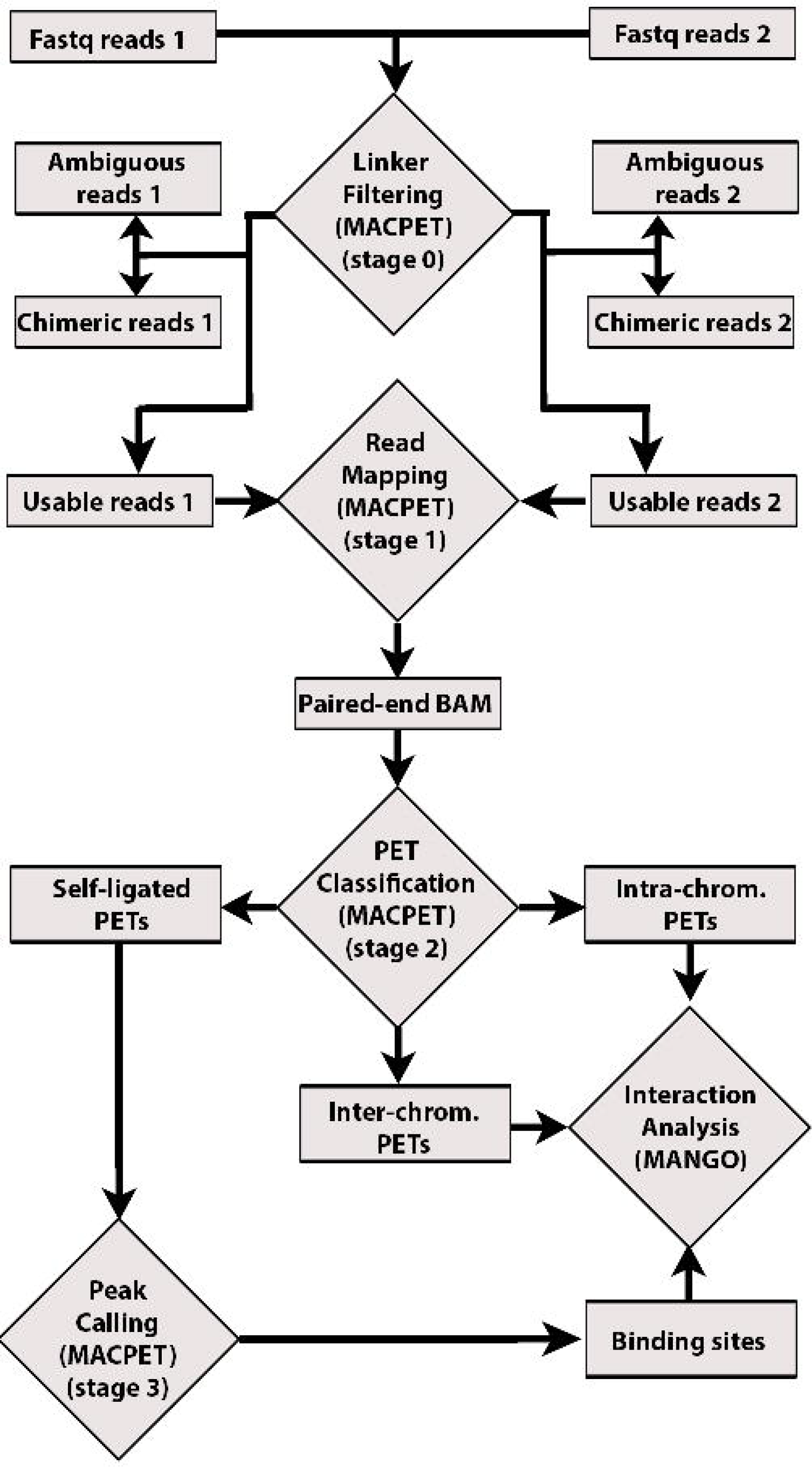
MACPET pipeline. Stage 0: MACPET takes the forward (1) and reverse (2) fastq files as input, as well as the user-specified barcode sequences for the half-linkers. It then classifies the PETs as ambiguous, chimeric and usable (non-chimeric). The half-linkers of the usable PETs are trimmed to release the two tags of each PET. Stage 1: The tags of the usable PETs are mapped separately to the reference genome and a paired-end BAM file is created. Stage 2: PETs are classified as self-ligated, intra-and inter-chromosomal. Stage 3: Self-ligated PETs are used for discovering PBSs. Finally, the PBSs as well as the intra-and inter-chromosomal PETs can be used in MANGO for interaction analysis.

### Stage 0: Linker filtering

MACPET identifies the half-linkers and classifies the PETs as ambiguous, chimeric and non-chimeric. It then removes the half-linker sequences from nonchimeric to reveal the two tags of each PET in the data. The trimmed non-chimeric PETs are used in the next stage of the analysis. This stage also removes PETs which include non-standard residues (for example the N letter).

### Stage 1: Mapping to the genome

MACPET maps the tags of the non-chimeric PETs separately to the reference genome using the Bowtie algorithm [19]. First the tags are mapped without allowing any mismatch and the uniquely mapped tags are kept. Then non-mapped tags are subject to a second run mapping with at most one mismatch and only the uniquely mapped tags are kept again. Note that this is the same process as the one proposed in [7]. PETs with both of their tags uniquely mapped with zero or one mismatch are used for constructing the paired-end BAM file which is used in subsequent stages. Finally, PETs with either tag overlapping any black listed regions of the corresponding genome are removed before continuing to the next stage of the analysis [20].

### Stage 2: PET classification

MACPET classifies the PETs into self-ligated, intra-and inter- chromosomal. Inter-chromosomal PETs can be easily separated as the tags of each PET are mapped on different chromosomes. For separating the other two categories, MACPET plots the histogram of the log-lengths of the PETs (using a bandwidth of 100 for each bin), spanning from the minimum to the maximum length of the PETs (see figure 2). The length of a PET is defined as the distance between its tags. It then applies the elbow method for finding a cut-off between the two populations. Thereafter, PETs for which both of their tags overlap with another PET’s tags (±1 bp) are removed and only one of those PETs is kept for reducing noise by PCR amplification procedures.

### Stage 3: Peak calling

At this stage MACPET uses only the self-ligated PETs for identifying candidate binding site locations. The genome is first segmented into non-overlapping regions, where each region has at least two overlapping self-ligated PETs. Each self-ligated PET in a region overlaps with at least one other self-ligated PET in the same region and no other self-ligated PET in any other region. In this way MACPET can analyse each region separately and ensure that a binding event can only belong to one region. An example of a region can be seen in figure 8.

**Figure 8.**
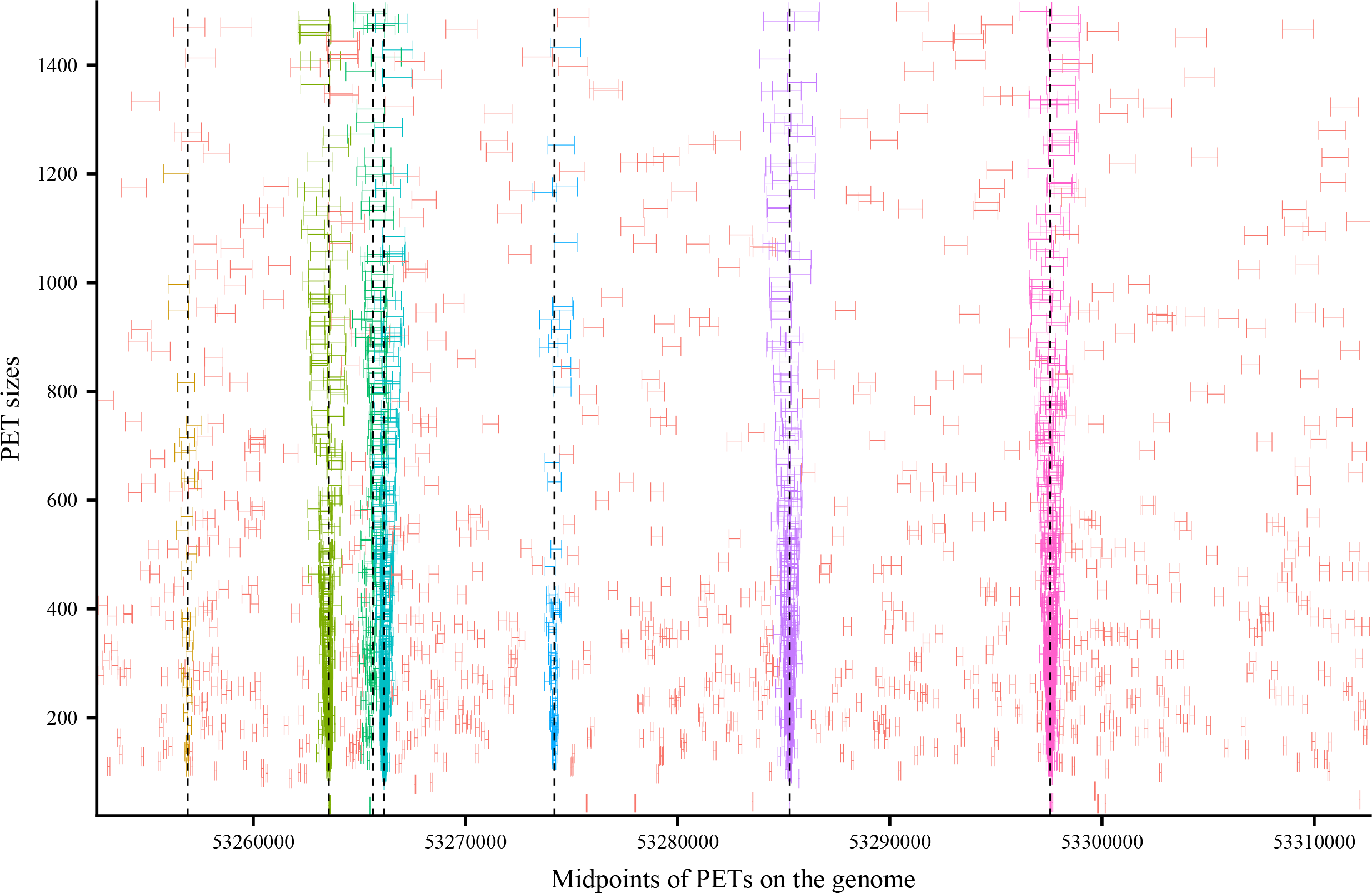
Illustration of a region. Illustration of a region in two dimensions. The x-axis is the midpoints of the PETs and the y-axis is the length of the PETs. Each segment represents a PET from its upstream to its downstream tag. Red colored PETs are classified as noise PETs by MACPET. The rest colors represent binding sites, where each color represents a different binding site. The dashed lines represent the exact binding location found by MACPET for each binding site.

### Distribution of tags in a Protein Binding Site

Self-ligated PETs which construct a PBS are products of the same type of protein which binds on approximately the same position across a set of identical genomes [7]. Therefore it should be reasonable to expect that self-ligated PETs in a specific PBS would have approximately the same characteristics regarding the positions of their upstream and downstream tags. On the other hand, noise PETs should have characteristics which differ from those in a PBS.

Consider a PBS *g* with *n*_*g*_ total self-ligated PETs and let 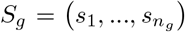 be the self-ligated PETs in that PBS, with *s*_*i*_ = (*x*_*i*_, *y*_*i*_) being the pair of upstream and downstream tags in self-ligated PET *i*, respectively. Although each tag of a self-ligated PET is mapped separately on the genome, with one tag not affecting the position of the other, we assume that *x*_*i*_ < *y*_i_. That is, we sort the tags of each self-ligated PET in increasing order for better representing the left and right stream tags. This of course creates a dependency between *x*_*i*_ and *y*_*i*_.

Because the PBS should have two peaks, one on each of its sides, MACPET models the left and right peaks of the PBS as a two-dimensional skew generalized t-distribution (SGT) distribution. The onedimensional SGT distribution is a five parameter distribution which models both skewness and long tails of the data [21]. It consists of three parameters 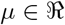, λ ∈ (−1, 1) and 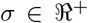 which represent the mode, skewness and scale, respectively, and two parameters *p,q* > 0 which are shape parameters. Here 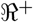 represents positive real numbers. It as been shown that for estimating the first three parameters, both of the shape parameters need to be known [21]. We choose *p* = 2 which leads to normal-type peak of the mode, and q = 1 which leads to heavy and long tails [21]. The one-dimensional SGT density function is: 
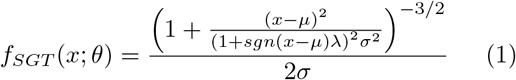
Where *θ* = (*μ*, λ *σ*), *sgn*(*x*) is the signum function which equals 1 if *x* > 0, −1 if *x* < 0 and if *x* = 0.

MACPET assumes that the two-dimensional density of a self-ligated PET *i*, *i* = 1,…, *n*_*g*_ in PBS *g* is *f*_*g*_(*s*_*i*_; *θ*_*g*_) = ⋀ (*θ*_*g*_) *f*_*xg*_(*x*_*i*_; *θ*_*xg*_)*f*_*yg*_(*y*_*i*_; *θ*_*yg*_) if *x*_*i*_ < *y*_*i*_ and 0 if *x*_*i*_ ≥ *y*_*i*_ Where *f*_*xg*_(*x*_*i*_; *θ*_*xg*_) and *f*_*yg*_(*y*_*i*_; *θ*_*yg*_) have the SGT density given in equation 1. Moreover, *θ*_*g*_ = (*θ*_*xg*_, *θ*_*yg*_) are the parameters of the PBS *g*, with *θ*_*xg*_ = (*μ*_*xg*_ λ_*xg*_, *σ*_*xg*_), and *θ*_*yg*_ = (*μ*_*yg*_ λ_*yg*_, *σ*_*yg*_) being
the parameters of the upstream and downstream peaks of the PBS respectively.

The term 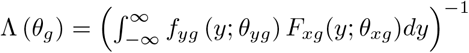 ensures that the function *f*_*g*_ (*s*_*i*_; *θ*_*g*_) integrates to 1, and hence is a valid probability density. Here *F*_*xg*_ is the cumulative distribution function of the SGT (see Section *Quantile function for the SGT distribution* in supplementary material available online).

MACPET models the left stream, 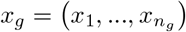, and right stream tags, 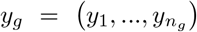, as negative and positive skewed towards the PBS location, respectively. This is achieved by imposing a hierarchical structure where λ_*xg*_ is restricted in the interval ( –1, 0] with the density function 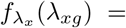 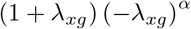, while λ_yg_ is restricted in the interval [0,1) with the density function 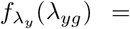 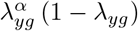. The value *α* = 39 has been chosen in order that λ_*xg*_ will tend towards −1, while λ_*yg*_ will tend towards +1.

Additionally, for ensuring that the left and right peaks are on their correct positions around the PBS, MACPET uses the reparametrization *μ*_*xg*_ = *μ*_*yg*_ – *k*_*g*_ where 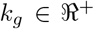. This ensures that the left peak of a PBS will always be on the left side of the right peak of the PBS on the genome.

The precise binding location is assumed to be between the two peak modes, that is (*μ*_*xg*_ + *μ*_*yg*_) /2. Furthermore, a 95% interval for the binding location is defined as (*Q*_*Lxg*_(0.05), *Q*_*Ryg*_(0.95)), where *Q*_*Lxg*_(0.05) is the 5% quantile of the upstream peak and *Q*_*Ryg*_(0.95) is the 95% quantile of the downstream peak (see Section *Quantile function for SGT distribution* in supplementary material available online).

### Modeling a region

Consider a region with *N* total self-ligated PETs and *G* total PBSs. Let *S* = (*s*_1_,…,*s*_*N*_) be the self-ligated PETs in that region, with *s*_*i*_ defined as before and *i* =1,…, *N*. MACPET models the region as a mixture of *G* clusters representing the PBSs and a noise cluster representing randomly distributed PETs in the region. That is, the density of S in the region is 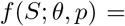 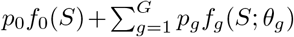, where *p* = (*p*_0_,…,*p*_*G*_) and *p*_*g*_ are the mixing probabilities of each cluster which sum to 1. Furthermore, *θ* = (*θ*_1_,…, *θ*_*G*_) and *g* = 1,…, *G* refer to the PBS clusters and *g* = 0 refers to the noise cluster. The noise cluster is assumed to be uniformly distributed with density *f*_0_(*x,y*) = 1/(2.5V) if *x* < *y* and 0 if *x* ≥ *y*, where *V* is the two-dimensional volume of the region. Note that the constant 2.5 increases the volume of the region and creates a slightly bigger area over the overlapping self-ligated PETs. By doing that, MACPET takes into account the noise level surrounding the region.

Taking into account the hierarchical structure for the λ parameters mentioned earlier in the text, the observed log-likelihood of the region is [22]:

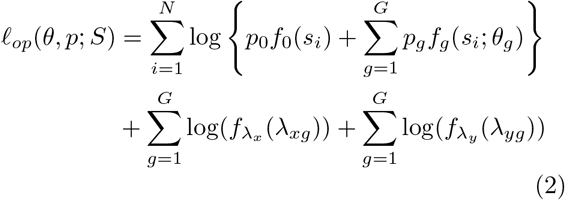

MACPET uses the Expectation/Conditional Maximization Either (ECME) algorithm for fitting the region model in equation 2 [23]. A detailed description of the estimation procedure can be found in Section *Deriving the likelihood for the estimation* in supplementary material available online.

### Inference

For assessing the significance of each candidate binding event, MACPET considers the quantile functions of the estimated candidate PBSs. Consider a candidate PBS *g* located at chromosome *C* and let (*Q*_*Lxg*_(0.05), *Q*_*Rxg*_(0.95)) and (*Q*_*Lyg*_(0.05), *Q*_*Ryg*_(0.95)) be the 95% confidence intervals for its upstream and downstream peaks, respectively (see Section *Quantile function for SGT distribution* in supplementary material available online). Furthermore, let *S*_*xg*_ and *S*_*yg*_ be the lengths of those intervals, and *N*_*Cx*_ and *N*_*Cy*_ be the total number of upstream and downstream tags on the chromosome *C*, respectively. Note that *N*_*Cx*_ = *N*_*Cy*_.

The null hypothesis for the upstream tags (*H*_*0xg*_) assumes that the number of tags in the upstream peak of *g* is random, following a Poisson distribution with in-tensity λ*xg* = *max*(2, λ_*Cx*_, λ_*wx*10_, λ_*wx*15_). Here λ_*Cx*_ — *N*_*Cx*_*S*_*xg*_/*S*_*C*_ is the expected number of upstream tags in the upstream peak, given the chromosome size *S*_*C*_. Furthermore, λ_*wx*10_ = *N*_*w*10*x*_*S*_*xg*_/(10*S*_*xg*_) and λ_*wx*15_ = *N*_*w*15*x*_*S*_*xg*_ /(15*S*_*xg*_) are the expected number of upstream tags in the upstream peak, by looking at a window of 10 and 15 times the size of the upstream peak, respectively. Furthermore, the constant 2 ensures that at least two tags have to exist in the peak interval in order to be considered significant. The analogous hypothesis is assumed for the downstream tags (*H*_0*yg*_) of *g*.

The null hypothesis for the candidate PBS *g* (*H*_0*g*_) assumes that *g* is not a PBS but a random sample of overlapping self-ligated PETs. Intuitively for *g* not being a true PBS, both of its upstream and downstream peaks need to be randomly formed, that is both *H*_0*xg*_ and *H*_0*yg*_ are valid. Let *E*_*g*_ be the event that *g* is not a true PBS and *E*_*xg*_ and *E*_*yg*_ the events under *H*_0*xg*_ and *H*_0*yg*_, respectively. Then under *H*_0*g*_ the upstream and downstream tags are assumed to be independent and thus *P*(*E*_*g*_) = *P*(*E*_*xg*_ ∩ *E*_*yg*_) = *P*(*E*_*xg*_)*P*(*E*_*yg*_). Therefore, the p-value for *g* can be defined as *p*_*g*_ = *p*_*xg*_*p*_*yg*_, where *p*_*xg*_ and *p*_*yg*_ are the p-values of the upstream and downstream peak, respectively. Finally, the p-values from all the PBSs are corrected using the Benjamini-Hochberg procedure [24].

Note that the quantile intervals are computed assuming ⋀ (*θ*_*g*_) = 1. That is, the quantile intervals are found using the marginal distributions of *x*_*g*_ and *y*_*g*_ under the assumption of independence between them. We use this assumption because computing the marginal distributions of *x*_*g*_ and *y*_*g*_, in case of dependence between them, was computationally intensive. This should not be a big violation of the model, however, as the estimated ⋀(*θ*_*g*_) value for the majority of the PBS in each dataset is very close to 1 (see figure S7 in supplementary material available online). There are a few values deviate a lot from 1, which could be the result of rounding errors while computing the ⋀ (*θ*_*g*_) integral. The reason that we still include the ⋀ (*θ*_*g*_) term when finding candidate PBS on the previous step of the algorithm is that we observed an increase in the speed of the algorithm as well as smoother convergence, while assuming that ⋀ (*θ*_*g*_) = 1 led to almost identical results but with slower speed.

## List of abbreviations

PBS: protein binding site; bp: base-pairs; PETs: paired-end tags; MACPET: Model-based Analysis for ChIA-PET; SGT: skew generalized t; ECME: Expectation/Conditional Maximization Either; FDR: false discovery rate;

### Ethics approval and consent to participate

Not applicable

### Consent for publication

Not applicable

### Availability of data and material

The MACPET algorithm is available on Bioconductor (https://bioconductor.org/packages/MACPET) under the public license GPL-3, and can be used on all platforms.

The datasets used during the current study are available in the NCBI repository [14]. More specifically:

ChIA-PET dataset for ESR1 TF from MCF-7 human cell line (GEO:GSM970212), available at (https://www.ncbi.nlm.nih.gov/geo/query/acc.cgi?acc=GSM970212).

ChIA-PET dataset for CTCF TF from MCF-7 human cell line (GEO:GSM970215), available at (https://www.ncbi.nlm.nih.gov/geo/query/acc.cgi?acc=GSM970215).

ChIA-PET dataset for CTCF TF from K562 human cell line (GEO:GSM970216), available at (https://www.ncbi.nlm.nih.gov/geo/query/acc.cgi?acc=GSM970216).

ChIA-PET dataset for histone H3K4me1 from K562 human cell line (GEO:GSM1436263), available at (https://www.ncbi.nlm.nih.gov/geo/query/acc.cgi?acc=GSM1436263).

ChIA-PET dataset for histone H3K27ac from K562 human cell line (GEO:GSM1436262), available at (https://www.ncbi.nlm.nih.gov/geo/query/acc.cgi?acc=GSM1436262).

ChIA-PET dataset for POL2 from K562 human cell line (GEO:GSM970213), available at (https://www.ncbi.nlm.nih.gov/geo/query/acc.cgi?acc=GSM970213).

All processed data are available at https://figshare.com/projects/MACPET_Model-based_Analysis_for_ChIA-PET/29473.

## Funding

Department of Mathematical Sciences, Norwegian University of Science and Technology, NTNU.

## Competing interests

The authors declare that they have no competing interests.

## Authors’ contributions

IV conceived the main idea behind MACPET, proved the statistical models, developed MACPET, performed the data analysis and drafted the manuscript. FD and MR proposed the project and helped to draft the manuscript. BL helped with the statistical models and the manuscript. All authors read and approved the final manuscript.

## Acknowledgements

The first author is grateful to Professor Giovanni Parmigiani for his kind help and great hospitality during this author’s stay at Dana Farber Cancer Institute in 2017.

## Additional Files

Additional file 1—Supplementary Material

Supplementary material which include figures from the rest of datasets as well as proofs and estimation procedure details can be found online.

---

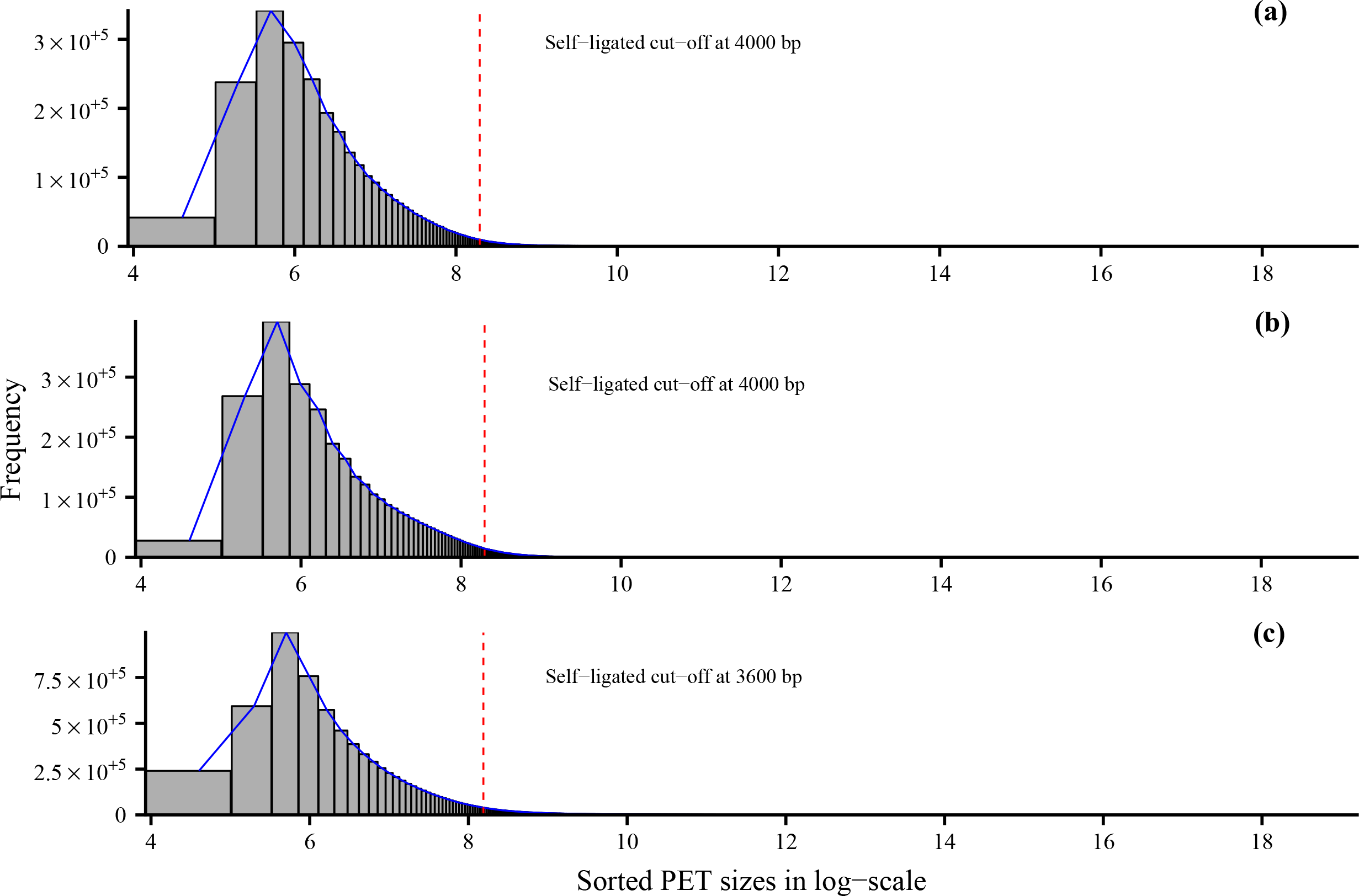

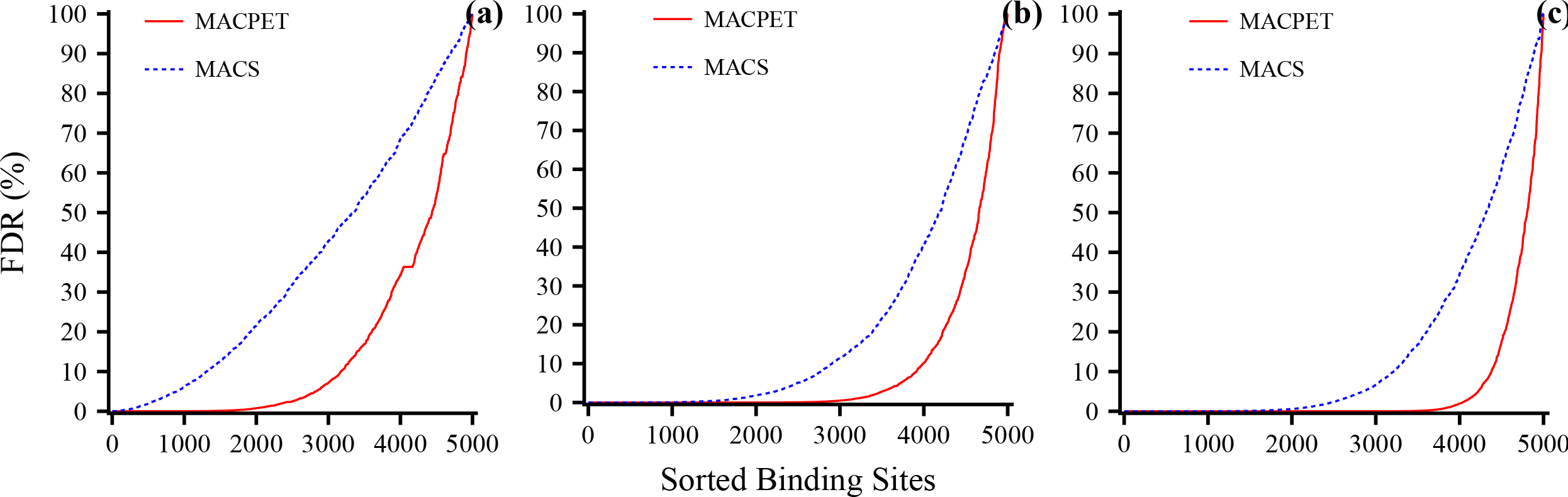

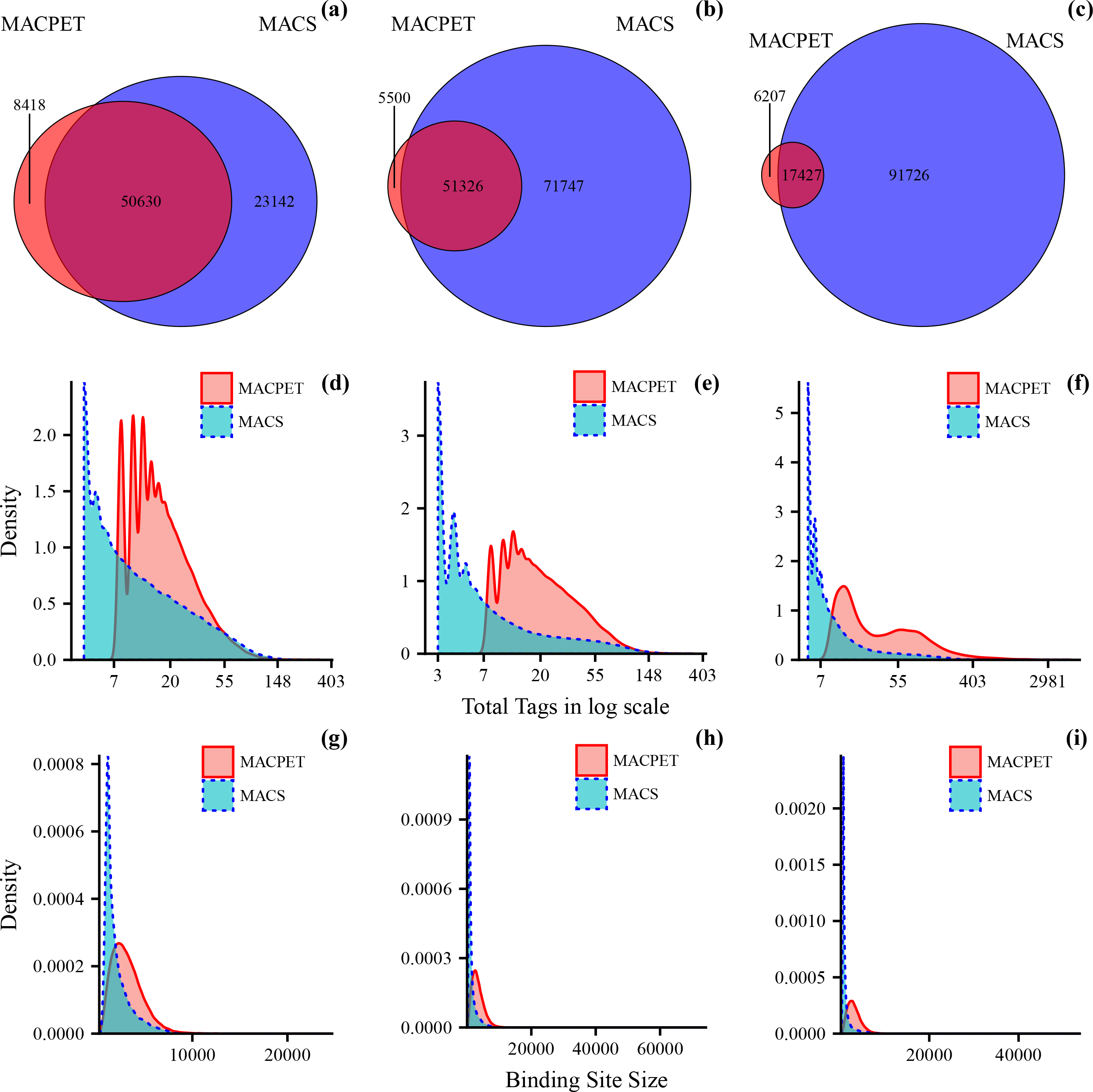

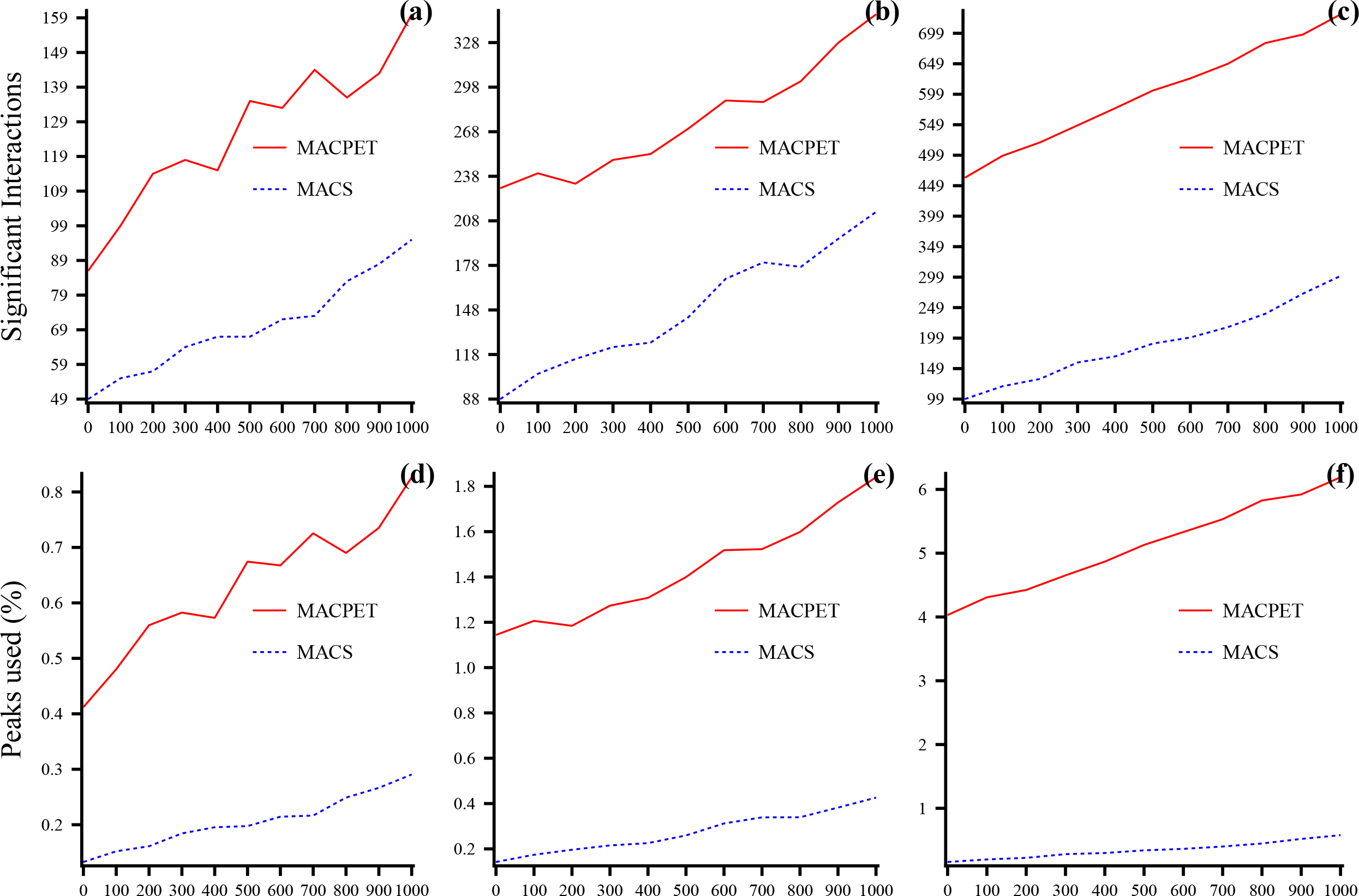

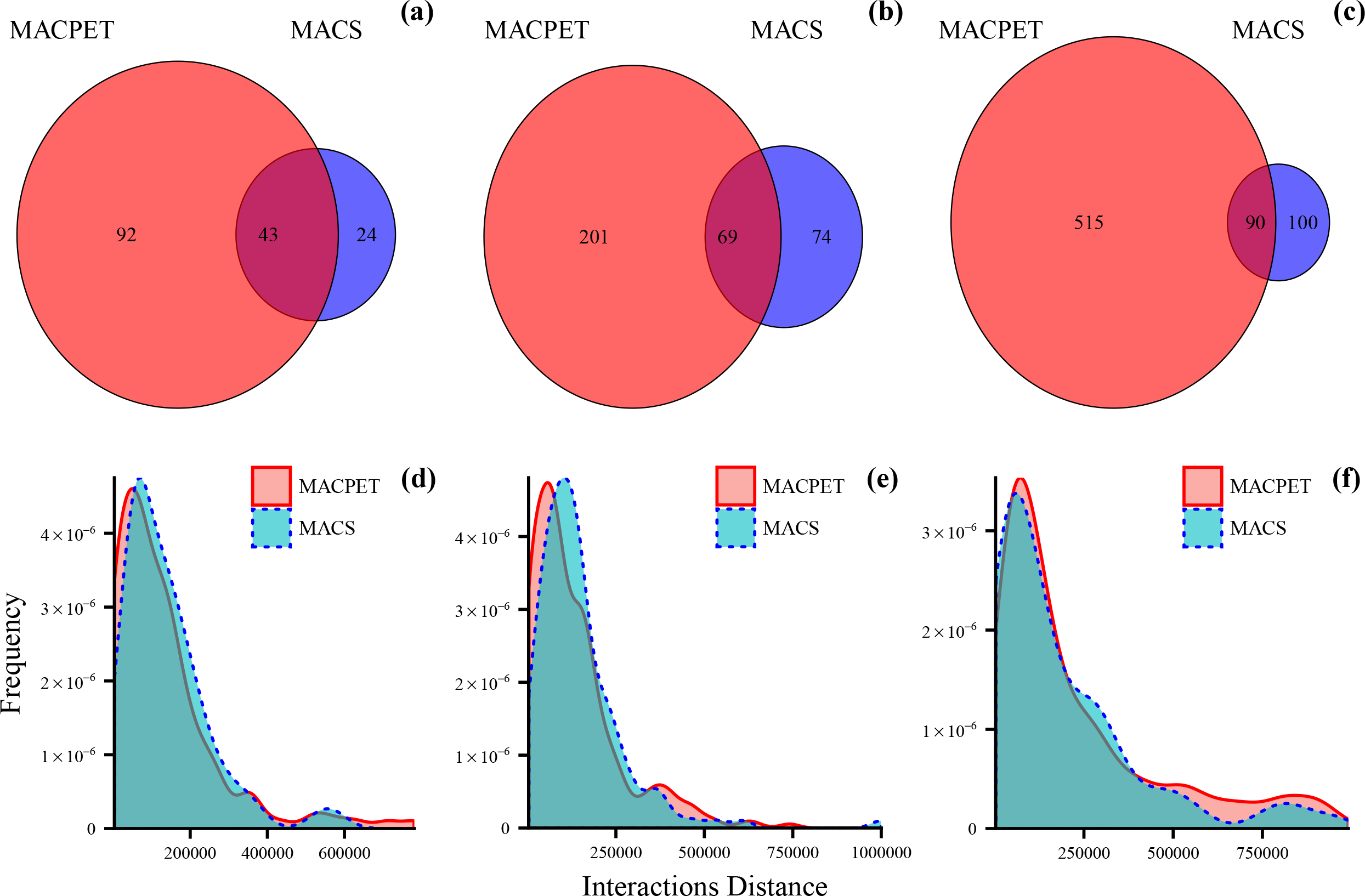

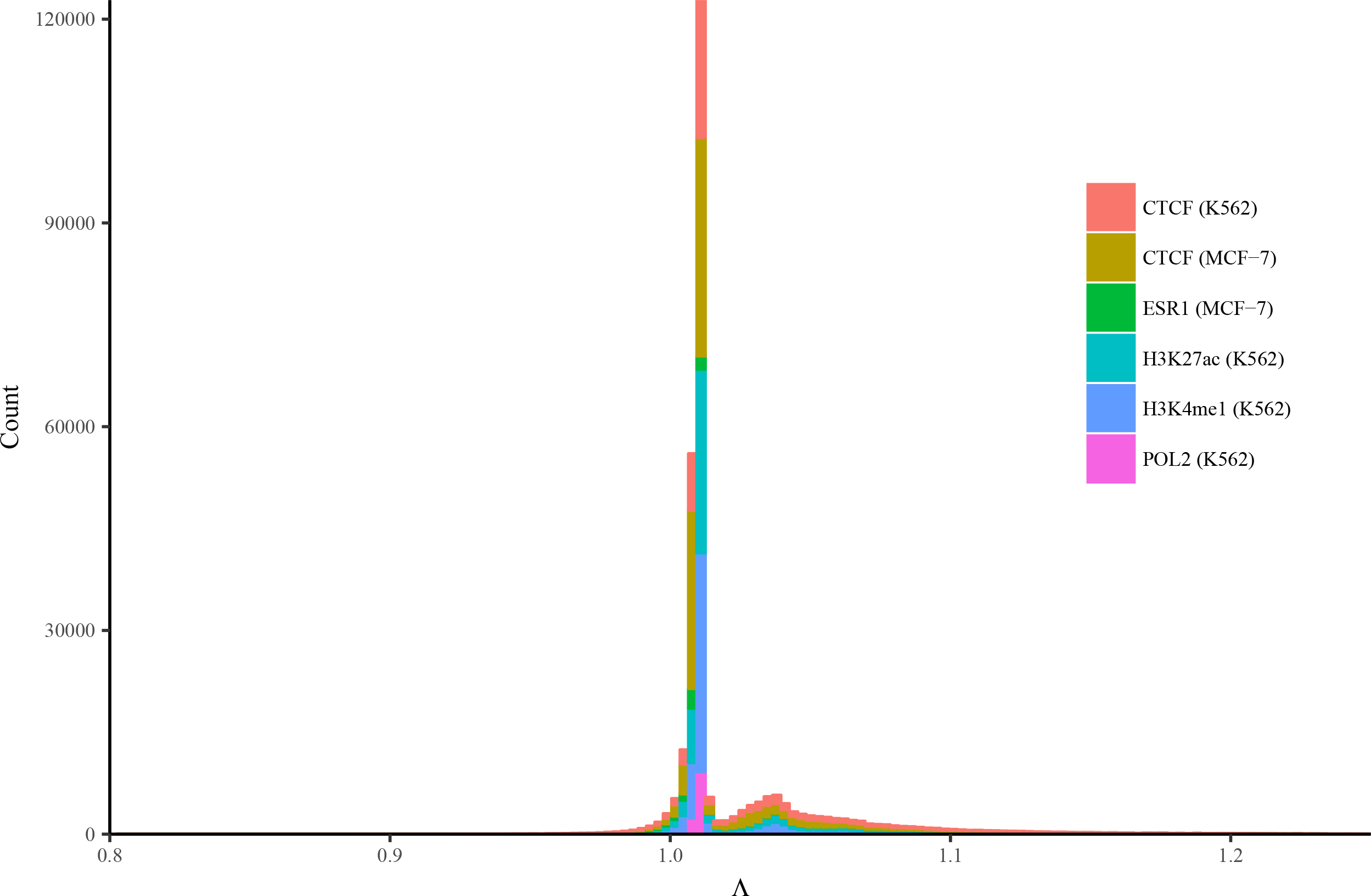

## References

1. Woodcock, C.L.: Chromatin architecture. Current Opinion in Structural Biology 16(2), 213–220 (2006). doi:10.1016/j.sbi.2006.02.005

2. Woodcock, C.L., Dimitrov, S.: Higher-order structure of chromatin and chromosomes. Current Opinion in Genetics and Development 11(2), 130–135 (2001). doi:10.1016/S0959-437X(00)00169-6

3. Dekker, J., Rippe, K., Dekker, M., Kleckner, N.: Capturing Chromosome Conformation. Science 295(5558), 1306–1311 (2002). doi:10.1126/science.1067799

4. Fraser, P., Bickmore, W.: Nuclear organization of the genome and the potential for gene regulation. Nature 447(7143), 413–417 (2007). doi:10.1038/nature05916

5. Wei, C.-L., Wu, Q., Vega, V.B., Chiu, K.P., Ng, P., Zhang, T., Shahab, A., Yong, H.C., Fu, Y., Weng, Z., Liu, J., Zhao, X.D., Chew, J.-L., Lee, Y.L., Kuznetsov, V.A., Sung, W.-K., Miller, L.D., Lim, B., Liu, E.T., Yu, Q., Ng, H.-H., Ruan, Y.: A global map of p53 transcription-factor binding sites in the human genome. Cell 124(1), 207–219 (2006). doi:10.1016/j.cell.2005.10.043

6. Fullwood, M.J., Liu, M.H., Pan, Y.F., Liu, J., Xu, H., Mohamed, Y.B., Orlov, Y.L., Velkov, S., Ho, A., Mei, P.H., Chew, E.G.Y., Huang, P.Y.H., Welboren, W.-J., Han, Y., Ooi, H.S., Ariyaratne, P.N., Vega, V.B., Luo, Y., Tan, P.Y., Choy, P.Y., Wansa, K.D.S.A., Zhao, B., Lim, K.S., Leow, S.C., Yow, J.S., Joseph, R., Li, H., Desai, K.V., Thomsen, J.S., Lee, Y.K., Karuturi, R.K.M., Herve, T., Bourque, G., Stunnenberg, H.G., Ruan, X., Cacheux-Rataboul, V., Sung, W.-K., Liu, E.T., Wei, C.-L., Cheung, E., Ruan, Y.: An oestrogen-receptor-[agr]-bound human chromatin interactome. Nature 462(7269), 58–64 (2009)

7. Li, G., Fullwood, M., Xu, H., Mulawadi, F.H., Velkov, S., Vega, V., Ariyaratne, P.N., Mohamed, Y.B., Ooi, H.-S., Tennakoon, C., Wei, C.-L., Ruan, Y., Sung, W.-K.: Chia-pet tool for comprehensive chromatin interaction analysis with paired-end tag sequencing. Genome Biology 11(2), 22 (2010). doi:10.1186/gb-2010-11-2-r22

8. Fullwood, M.J., Wei, C.-L., Liu, E.T., Ruan, Y.: Next-generation dna sequencing of paired-end tags (pet) for transcriptome and genome analyses. Genome Research 19(4), 521–532 (2009). doi:10.1101/gr.074906.107. http://genome.cshlp.org/content/19/4/521.full.pdf+html

9. Chiu, K., Wong, C.-H., Chen, Q., Ariyaratne, P., Ooi, H., Wei, C.-L., Sung, W.-K., Ruan, Y.: Pet-tool: a software suite for comprehensive processing and managing of paired-end ditag (pet) sequence data. BMC Bioinformatics 7(1), 390 (2006). doi:10.1186/1471-2105-7-390

10. Harbers, M., Kahl, G.: Tag-based Next Generation Sequencing, pp. 186–209. Wiley-Blackwell, 111 River Street, New Jersey, NJ 07030-5774, USA (2012). Chap. 12

11. Do, K.-A., Qin, Z.S., Vannucci, M. (eds.): Advances in Statistical Bioinformatics, pp. 138–169. Cambridge University Press, 1 Liberty Plaza, Floor 20, New York, NY 10006, USA (2013). Chap. 7. Cambridge Books Online. http://dx.doi.org/10.1017/CBO9781139226448

12. Zhang, Y., Liu, T., Meyer, C., Eeckhoute, J., Johnson, D., Bernstein, B., Nusbaum, C., Myers, R., Brown, M., Li, W., Liu, X.S.: Model-based analysis of chip-seq (macs). Genome Biology 9(9), 137 (2008). doi:10.1186/gb-2008-9-9-r137

13. Phanstiel, D.H., Boyle, A.P., Heidari, N., Snyder, M.P.: Mango: a bias-correcting chia-pet analysis pipeline. Bioinformatics 31(19), 3092–3098 (2015). doi:10.1093/bioinformatics/btv336

14. NCBI Resource Coordinators: Database resources of the national center for biotechnology information. Nucleic Acids Research 44(Database issue), 7–19 (2016). doi:10.1093/nar/gkv1290

15. Droit, A., Gottardo, R., Robertson, G., Li, L.: rgadem: de novo motif discovery. r package version 2.20.0. (2014)

16. Mercier, E., Gottardo, R.: Motiv: Motif identification and validation. r package version 1.28.0. (2014)

17. Li, L.: gadem: A genetic algorithm guided formation of spaced dyads coupled with an em algorithm for motif discovery. Journal of Computational Biology 16(2), 317–329 (2009). doi:10.1089/cmb.2008.16TT

18. Zhang, X., Robertson, G., Krzywinski, M., Ning, K., Droit, A., Jones, S., Gottardo, R.: Pics: Probabilistic inference for chip-seq. Biometrics 67(1), 151–163 (2011). doi:10.1111/j.1541-0420.2010.01441.x

19. Langmead, B., Trapnell, C., Pop, M., Salzberg, S.L.: Ultrafast and memory-efficient alignment of short dna sequences to the human genome. Genome Biology 10(3), 25 (2009). doi:10.1186/gb-2009-10-3-r25

20. ENCODE Project Consortium: An integrated encyclopedia of dna elements in the human genome. Nature 489(7414), 57–74 (2012)

21. Arslan, O., Genç,A.i.: The skew generalized t distribution as the scale mixture of a skew exponential power distribution and its applications in robust estimation. Statistics 43(5), 481–498 (2009). doi:10.1080/02331880802401241. http://dx.doi.org/10.1080/02331880802401241

22. Fraley, C., Raftery, A.E.: Bayesian regularization for normal mixture estimation and model-based clustering. J. Classif. 24(2), 155–181 (2007). doi:10.1007/s00357-007-0004-5

23. Liu, C., Rubin, D.B.: The ecme algorithm: A simple extension of em and ecm with faster monotone convergence. Biometrika 81(4), 633–648 (1994)

24. Benjamini, Y., Hochberg, Y.: Controlling the false discovery rate: A practical and powerful approach to multiple testing. Journal of the Royal Statistical Society. Series B (Methodological) 57(1), 289–300 (1995)

